# TNFR1/p38αMAPK signaling in Nex+ supraspinal neurons regulates sex-specific chronic neuropathic pain

**DOI:** 10.1101/2023.08.01.551503

**Authors:** Kathryn A. Swanson, Kayla L. Nguyen, Shruti Gupta, Jerome Ricard, John R. Bethea

## Abstract

Upregulation of soluble tumor necrosis factor (sTNF) cytokine signaling through TNF receptor 1 (TNFR1) and subsequent neuronal hyperexcitability are observed in both animal models and human chronic neuropathic pain (CNP) (Clark et al., 2013; Empl et al., 2001; Ji et al., 2018; Lindenlaub and Sommer, 2003). To test the hypothesis that supraspinal circuitry is critical to pain chronification, we studied the intersect between supraspinal TNFR1 mediated neuronal signaling and sex specificity by selectively removing TNFR1 in Nex+ neurons in adult mice (NexCre^ERT2^::TNFR1^f/f^). We determined that following chronic constriction injury (CCI), pain resolves in males; however, female acute pain transitions to chronic. Subsequently, we investigated two downstream pathways, p38MAPK and NF-κB, important in TNFR1 signaling and injury response. We detected p38αMAPK and NF-κB activation in male cortical tissue; however, p38αMAPK phosphorylation was reduced in NexCre^ERT2^::TNFR1^f/f^ males. We observed similar behavioral results following CCI in NexCre^ERT2^::p38αMAPK^f/f^ mice. Previously, we established estrogen’s ability to modulate sTNF/TNFR1 signaling in CNP, which may contribute to female prevalence of CNP (Bouhassira et al., 2008; Claiborne et al., 2006; de Mos et al., 2007; Del Rivero et al., 2019; Li et al., 2009). To explore the intersection between estrogen and inflammation in CNP we used a combination therapy of an estrogen receptor β (ER β) inhibitor with a sTNF/TNFR1 or general p38MAPK inhibitor. We determined both combination therapies lend “male-like” therapeutic relief to females following CCI. These data suggest that TNFR1/p38αMAPK signaling in Nex+ neurons in CNP is male-specific and lack of therapeutic efficacy following sTNF inhibition in females is due to ER β interference. These studies highlight sex-specific differences in pathways important to pain chronification and elucidate potential therapeutic strategies that would be effective in both sexes.

## 1. INTRODUCTION

Chronic neuropathic pain (CNP) afflicts 7-20% of adults and can result from disease or injury to the somatosensory nervous system (Clark et al., 2013; Millecamps et al., 2023; Nahin, 2015). Critical mediators of persistent pain include neuroinflammation, such as increased levels of the inflammatory cytokine soluble tumor necrosis factor (sTNF), and maladaptive plasticity within the central nervous system (CNS) which extend beyond resolution of the initial insult (Costigan et al., 2009). While 7-20% of adults suffer from CNP, there is greater prevalence seen in females Sorge and Totsch, 2017). Previous research has established the influence of biochemical sex differences that contribute to this health disparity and established that sex hormones are linked to pain tolerance (Claiborne et al., 2006; Li et al., 2009). Specifically, estradiol-17β has been linked to enhanced immune reactivity, but it remains unclear whether it serves in a protective or injurious capacity for pain development (Deng et al., 2017; Lee et al., 2018; Vacca et al., 2016). However, the intersection between neuroinflammation, maladaptive plasticity, and estrogen remains unclear.

In several animal models of CNP, increased production of sTNF is observed at both acute and chronic timepoints following injury (Bethea et al., 1999; Del Rivero et al., 2019; Ruuls et al., 2001; Zhang et al., 2013). sTNF preferentially signals through its receptor, TNFR1, which is ubiquitously expressed. Under physiological conditions, sTNF/TNFR1 signaling regulates glutamatergic receptor transmission and mediates glial production of pro-inflammatory cytokines (Bradley, 2008; Stellwagen, 2005; Woods et al., 2021). However, under pathological conditions TNFR1 signaling is associated with both initiation and maintenance of neuroinflammation in part through activation of mitogen-activated protein kinase (MAPK) transduction pathways (Sabio and Davis, 2014; Yang et al., 2018). Interestingly, increased production of sTNF is not limited to the site of injury. After chronic constriction injury (CCI) of the sciatic nerve increased expression of sTNF is observed in both the peripheral and central nervous systems (Andrade et al., 2016). Studies in human patients reflect similar dispersion of sTNF levels where pain following a peripheral surgery corresponded with increased central sTNF levels (Andrade et al., 2016). This may be a result of downstream effects resulting from increased sTNF/TNFR1 activity.

Specifically, increased sTNF/TNFR1 signaling is associated with increased phosphorylation of p38MAPK which is linked to chronic pain states (Jin et al., 2003; Schäfers et al., 2003; Tsuda et al., 2004). Increased MAPK signaling then further stimulates sTNF production by the target cell creating a positive feedback loop exacerbating cytokine expression levels (Sabio and Davis, 2014). p38MAPK phosphorylation is a known indicator of proinflammatory microglia and may promote prolonged inflammation in the spinal cord. Increased sTNF levels have been directly associated with increased activation of proinflammatory microglia both in the spinal cord and dorsal root ganglia of several models, including CCI (Zhang et al., 2013). Together, these data would suggest that increased production of sTNF may trigger proinflammatory microglial activity via p38MAPK phosphorylation.

A strong correlation exists between the presence of CNS maladaptive plasticity, highlighted by increased excitatory signaling, and CNP. Specifically, increased production of sTNF by proinflammatory glia can cause nearby neurons to overexpress membranous glutamate receptors Haroon et al., 2017; Parsons and Raymond, 2014). Overexcitation promotes rapid synaptic remodeling characterized by increased expression of excitatory receptors, such as AMPAR and NMDAR. Supraspinal structures undergoing such adverse changes have been linked to aversive learning and memory outcomes (Baliki and Apkarian, 2015; Mansour et al., 2014; Yang and Chang, 2019). While both microglia and neurons respond and contribute to maladaptive CNS changes, their role in CNP remains unclear.

In this study, we find that chronic pain in males is dependent on TNFR1 signaling and p38αMAPK activation in a subset of supraspinal Nex+ excitatory neurons, sometimes referred to as neuronal differentiation 6 (NeuroD6) neurons. However, when estrogen receptor β (ER β) is inhibited, females become more ‘male-like’ with respect to the role of sTNF in the development and treatment of chronic pain.

## 2. MATERIALS AND METHODS

### 2.1 Mice

Male and female mice were bred in-house to generate NexCre^ERT2^::TNFR1^f/f^ and NexCre^ERT2^::p38αMAPK^loxP/loxP^ by crossing Neurod6^tm2.1(cre/ERT2)Kan^ mutants (MGI: 5308766/ Dr. K. Nave) with either TNFR1^loxP/loxP^ controls (EMMA 07099/ Dr. G. Kollias) or Mapk14^tm1.2Lex/YiwaJ^ mutants (Jackson Lab, #031129). Littermate controls were used for all studies involving in-house bred mice. For drug studies, 10-week-old C57Bl/6J female and male mice were obtained from Jackson Lab (#000664). Animals were housed in a 12/12 h light/dark cycle room and given food and water *ad libitum*. Mice were acclimated to the colony housing room for 2 weeks prior to experiments. All animal-use experimental protocols were carried out with approval of Drexel University’s Institutional Animal Care and Use Committee (#20295, #20895) and in accordance with the United States Public Health Service’s Policy on Humane Care and Use of Laboratory Animals.

### 2.2 Tamoxifen injections

To induce Cre recombination, tamoxifen (Sigma Aldrich, T5684; 75mg/kg) was administered via intraperitoneal injection for 5 consecutive days to both mutants and floxed controls at 10-weeks-old. Tamoxifen administration was completed 2 weeks prior to surgery to ensure recombination efficacy.

### 2.3 Chronic Constriction Injury (CCI)

As previously described by del Rivero et al, we performed chronic constriction injury (CCI) of the right sciatic nerve in 12-week-old mice. Prior to surgery, anesthesia was provided via intraperitoneal injection of ketamine (100 mg/kg) and xylazine (10 mg/kg). The sciatic nerve was exposed with an incision made in the mid-thigh. Once exposed, the sciatic nerve was tied with 3 silk ligatures (Oasis Silk 6-0, MV-711-V), spaced 1mm-1.5mm apart starting 3mm proximal to the sciatic bifurcation, and each subsequent ligature placed closer to the hip. The overlying skin was then closed with nylon sutures (Oasis Nylon 5-0, MV-661).

### 2.4 Behavioral Testing

The von Frey test was performed 3 days prior to CCI, establishing baseline mechanical sensitivity, and repeated weekly post-CCI. To perform von Frey testing, mice were placed in clear individual boxes on a wire mesh and habituated for 40 minutes prior to testing. Using an up-down method, an incremental series of 8 von Frey filaments (force range: .02g-2g) were applied to the plantar surface of the hind paw, as previously described (Chaplan et al., 1994). For all mice, we tested both ipsilateral and contralateral hind paws. Each test consisted of 6 trials, with 5 minutes between each trial. Positive responses to a filament were noted as both hind paw withdrawal and a cognitive component, such as guarding behavior or hind paw grooming. This dual attention/response criteria allows the experimenter to discount reflexive movement as a false-positive response. We quantified responses according to the standard up-down analysis method (26). For behavioral testing, the contralateral hind paw served as the uninjured side and the ipsilateral hind paw served as the injured side. The experimenter remained blinded to experimental conditions/treatments until the termination of the experiment.

### 2.5 Drugs

Faslodex (20 mg/kg; Abcam ab120131), a clinically used general estrogen receptor inhibitor, and ERβ specific inhibitor PHTPP (1.7mg/kg; Sigma Aldrich SML1355) were diluted in corn oil and administered systemically. Drug or vehicle (corn oil) was administered via intraperitoneal injection first at CCI. Treatment with Xpro1595 (10 mg/kg; intraperitoneal; INmuneBio), a dominant negative sTNF inhibitor, or vehicle (0.1M PBS) began 7 days post-CCI. Previously mentioned drugs were given every 48 hours until experiment termination. SB203580, a p38_αβƴ_ MAPK inhibitor (Invivogen tlrl-sb20), was diluted to 4.54mg/mL in PBS and delivered to the CNS at 0.11µL/hr via osmotic pump (Azlet 1004) administering the drug at 0.5µg/hr with intrathecal cannula (Azlet Brain Infusion Kit3). Pumps containing SB203580 or vehicle were inserted during CCI via cannulation at the T12 vertebrae level.

### 2.6 Tissue Collection

#### 2.6a Whole tissue protein extract

At 6 weeks post-injury, mice were anesthetized with xylazine (43mg/kg) and ketamine (215mg/kg) and perfused with 0.1M PBS. Tissue was isolated from the sensorimotor cortex, hippocampus, and lower thoracic and lumbar spinal cord were removed and frozen on dry ice. Tissue was homogenized in RIPA buffer (10 mM Tris-HCl pH 7.4, 1 mM EDTA, 0.5 mM EGTA, 1% NP-40, 0.1% Na Deoxycholate, 0.1% SDS, 140 mM NaCl) with protease and phosphatase inhibitors (Santa Cruz and Biovision, respectively). After centrifugation (14000 rpm, 4°C, 15 minutes) the supernatant was isolated and stored at −80°C.

#### 2.6b Synaptosome protein extract

For synaptic preparations, mice were perfused with HEPES buffer (2mM HEPES, 0.32 M sucrose, 10 mM sodium pyrophosphate, 10 mM sodium fluoride, 2 mM EGTA, 1 mM sodium orthovanadate, and 0.1 mM PMSF). Cortical tissue was isolated, frozen on dry ice and homogenized in HEPES buffer with 1 µg/mL leupeptin and 1 µg/mL aprotinin. First, the homogenate was centrifuged (3500 rpm, 4°C, 10 minutes) and the resulting supernatant was removed, and the pellet was kept as a purity control. The supernatant was centrifuged (13500 rpm, 4°C, 15 minutes) and the remaining pellet, which is the synaptosomal fraction, was resuspended in HEPES buffer while the supernatant was kept also for purity control.

### 2.7 Western Blot

Protein extracts were run on sodium dodecyl sulphate (SDS)-polyacrylamide gels and transferred to nitrocellulose membranes (Turbo blot, Biorad). Membranes were blocked for 2 hours in 5% non-fat milk (BioRad) in 1x TBS-T (10 mM tris-HCl, pH 7.5, 150 mM NaCl, 0.1% Tween-20). Primary antibodies were diluted in blocking solution and incubated at 4°C overnight. Membranes were probed with antibodies against: estrogen receptor β (Erβ, rabbit, 1:1000, Invitrogen PA1-311), tumor necrosis factor receptor 1 (TNFR1, rabbit, 1:1000, Cell Signaling 13377), N-methyl-D-aspartate receptor 1 (NMDAR1, rabbit, 1:1000, Invitrogen PA5-34599), N-methyl-D-aspartate receptor 2B (NMDAR2B, rabbit, 1:1000, Cell Signaling 4270), nuclear factor kappa B (NFκB, rabbit, 1:1000, Abcam ab16502), phosphorylated nuclear factor kappa B (phospho-NFκB, rabbit, 1:1000, Abcam ab76302), p38 mitogen-activated protein kinase (p38MAPK, mouse, 1:2000, Biolegend 602651), and phosphorylated-p38MAPK (phospho-p38MAPK, rabbit, 1:3000, Cell Signaling 4511). All primary antibodies were followed by horseradish peroxidase-conjugated species-specific secondary antibodies. Proteins were visualized with a chemiluminescent substrate (West Pico, ThermoFischer Scientific) and band intensity was quantified using FIJI software (Rasband W, NIH). Proteins were normalized to total protein per sample using Ponceau S staining solution (ThermoFisher) and subsequently normalized to the control conditions for graphing quantification.

### 2.8 Immunohistochemistry

Mice were transcardially perfused with 4% paraformaldehyde (Sigma-Aldrich) in PBS. Dissected brains were post-fixed in 4% PFA overnight and subsequently dehydrated in 25% sucrose for 48 hours at 4 °C. Tissue was frozen in OCT (Tissue-Tek) and stored at −80°C until sectioning. Brains were sectioned at 40 µm and preserved as free-floating sections in 20 mM phosphate buffer (20% glycerol/0.01% sodium azide) at 4 °C. Sections were washed three times in 0.1M PBS at room temperature (10 min/wash) and incubated in sodium citrate buffer (pH 6.8) at 60°C for 30 minutes for antigen retrieval. Washes were repeated and sections were then placed in blocking solution (10% NGS/0.3% Triton-X/0.1M PBS) for two hours at room temperature Sections were incubated in the following primary antibodies overnight at 4 °C: ERβ (rabbit, 1:200, Invitrogen PA1-311) and NeuN (mouse, 1:100, Millipore MAB337).Washes were repeated and the following secondary antibodies were applied for 30 minutes at room temperature: AlexaFluor594 (goat anti-rabbit, 1:500, Thermo Fisher Scientific A-11012) and AlexaFluor488 (goat anti-mouse, 1:500, Thermo Fisher Scientific A-11029). Sections were washed, nuclei were stained with Hoechst 33258 (1:10000, Anaspec AS-83219), and then slides were cover-slipped (Fluoro-Gel, Electron Microscopy Sciences).

### 2.9 Statistics

Data was analyzed and graphed on GraphPad Prism software (version 9.4.1). Western blots comparing 3 or more groups were analyzed by one-way ANOVA (α=0.05), with each group compared to each other using Tukey’s multiple comparisons test. Western blots comparing 2 groups were analyzed using a non-parametric, two-tailed Mann-Whitney t-test (α=0.05). Changes in mechanical thresholds were evaluated by two-way ANOVA (α=0.05), with time post-CCI and experimental group (genotype or treatment) as factors. The Geisser-Greenhouse correction was used to control for within group variability following by Tukey’s multiple comparisons test for all timepoints. Values for all data are expressed as mean ± SEM. No data outliers were identified (ROUT test, Q=1%).

## 3. RESULTS

### 3.1 TNFR1 signaling in supraspinal excitatory projection neurons is required for CNP in males, not females

Previous work from our group demonstrated sex differences from systemic disruption of sTNF/TNFR1 signaling; specifically global TNFR1 knockout prevented post-CCI allodynia in males while females had only partial alleviation (Del Rivero et al., 2019). Based on these data, we hypothesized that supraspinal TNFR1 signaling is critical for processing somatosensory inputs that then drive CNP. To test this, we generated NexCre^ERT2^::TNFR1^f/f^ mice to interrogate the development of CNP in males and females. Nex, a neuronal basic helix-loop-helix protein, is expressed among supraspinal glutamatergic pyramidal projection neurons which project processes between the neocortex and hippocampus (Agarwal et al., 2012). NexCre^ERT2^ mice are an inducible Cre line that specifically targets cortical and hippocampal pyramidal neurons that project between both regions (Agarwal et al., 2012). By selectively ablating TNFR1, which is ubiquitously expressed, among the Nex+ neuronal subset, we are able determine the contribution of selective supraspinal TNFR1 signaling in excitatory neurons. TNFR1 selective deletion in these neurons prevents the development of chronic pain solely in males and no effect is observed on acute pain for either sex (**Fig. 1A, B**). To ensure recombination efficiency, we measured TNFR1 expression levels in the cortex (Cx), hippocampus (Hc), and spinal cord (SC). We observed a significant reduction in cortical and hippocampal TNFR1 expression levels for both males and females to the extent anticipated for TNFR1 loss in the Nex+ neuronal subset; however, no changes were observed in the spinal cord (**Fig. 1C-H**).

**Figure 1.**
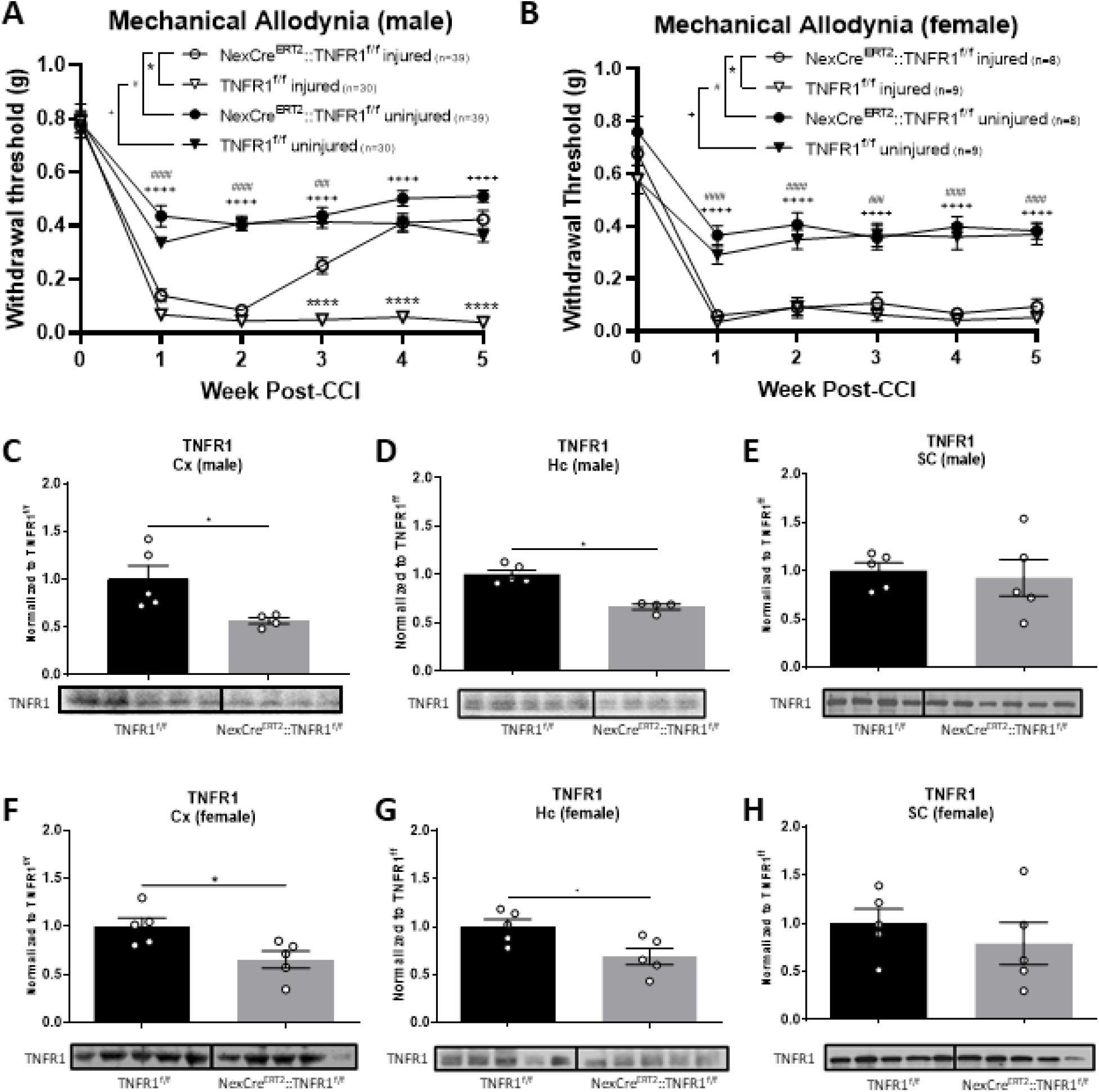
Chronic pain in males is dependent on TNFR1 signaling in supraspinal excitatory projection neurons. Withdrawal thresholds were measured over 5 weeks post-CCI and we observed that mechanical allodynia is reduced in NexCre::TNFR1^f/f^ (A) males, but not (B) females. Naive male and female TNFR1 expression in NexCre::TNFR1^f/f^ mice is reduced in sensorimotor cortex (C,F) and hippocampal (D,G) tissue, but no changes were observed in the lumbar spinal cord (E,H) (*p<0.05, **p<0.01, *** p<0.001, **** p<0.0001; Data represent mean ± SEM).

### 3.2 p38MAPK signaling in supraspinal excitatory projection neurons is required for CNP in males, not females

TNFR1 signaling is known to elicit downstream effects among both NF-κB and p38MAPK signaling pathways. Pharmacological inhibition of p38MAPK has been demonstrated to be therapeutic for CNP in a sexually dimorphic manner (Luo et al., 2018; Sorge et al., 2015; Taves et al., 2016a). Mechanistically, this has largely been attributed to attenuating p38MAPK-dependent microglial activation in the spinal cord of male, but not female mice (Taves et al., 2016b). However, following injury we observed a significant reduction in cortical p38αMAPK activation selectively in male NexCre^ERT2^::TNFR1^f/f^ mice (**Fig. 2B, E)**. Cortical NF-κB activation in males and females remained elevated in NexCre^ERT2^::TNFR1^f/f^ mice following injury compared to naïve controls (**Fig. 2A, D**). To determine whether p38αMAPK activation in these neurons is critical for development of chronic pain we generated NexCre^ERT2^::p38αMAPK^f/f^ mice. As shown in **Fig. 2C, F**, selective deletion of p38αMAPK in supraspinal Nex+ neurons prevented the development of chronic but not acute pain in male, but not female mice. To our knowledge, this is the first genetic evidence that p38αMAPK signaling in supraspinal excitatory projection neurons is crucial for the sex-specific effects of CNP. Whole tissue western blots of total p38αMAPK confirmed successful recombination in males and females with decreased cortical (**supplemental Fig. 2A,B**), but not spinal (**supplemental Fig. 2C,D**), p38αMAPK activation.

**Figure 2.**
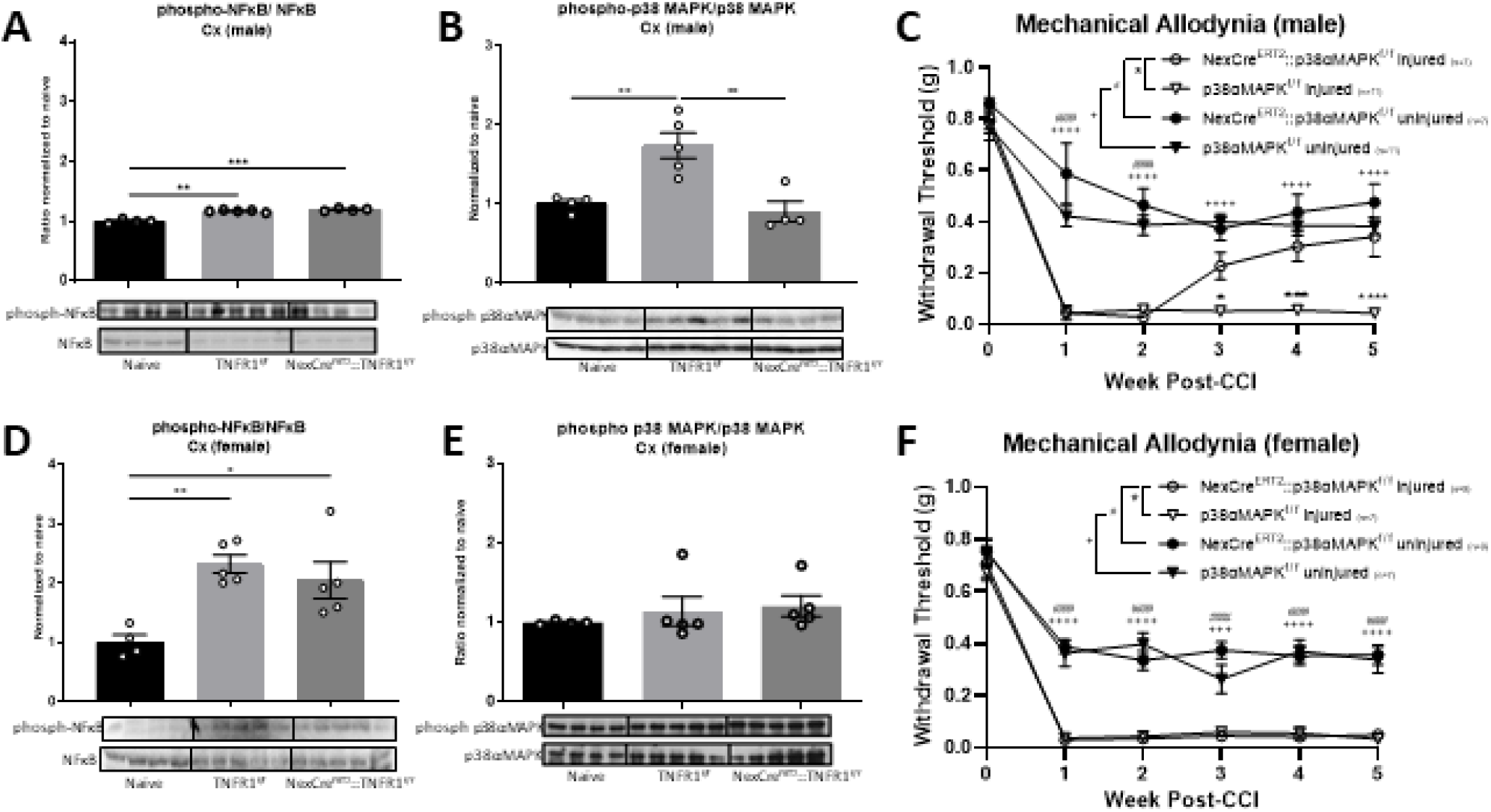
Cortical activation of p38αMAPK following injury is reliant on supraspinal excitatory projection neurons in males. Cortical p65-NFκB activation remains elevated following injury in NexCre::TNFR1^f/f^ (A) males and (D) females. (B) p38αMAPK activation in males increases following injury and is attenuated in the NexCre::TNFR1^f/f^ mice, but (E) no changes are observed compared to female controls. p38αMAPK deletion in Nex+ neurons attenuates mechanical allodynia in (C) males, but not (F) females (*p<0.05, **p<0.01, *** p<0.001, **** p<0.0001; Data represent mean ± SEM).

### 3.3 sTNF/TNFR1, maladaptive plasticity and chronic pain

Maladaptive plasticity within cortical areas, such as the primary somatosensory cortex (S1), has been demonstrated to play an important role in the development of CNP. In humans, the extent of pathological plasticity in S1 appears to be directly related to the sensorimotor intensity of chronic pain following peripheral or central injuries (Kim et al., 2017; Moseley and Flor, 2012; Wrigley et al., 2009). Importantly, attenuation of maladaptive neuronal plasticity is associated with reductions in CNP (Bak et al., 2021). Unfortunately, the intersection of maladaptive plasticity, neuroinflammation and chronic pain has not been adequately investigated. To address this, we investigated changes in glutamate receptor expression in cortex of naïve, TNFR1^f/f^ and NexCre^ERT2^::TNFR1^f/f^ mice. We observed a robust increase in NMDAR1 expression in males following CCI in TNFR1^f/f^ controls, but an attenuation in the NexCre^ERT2^::TNFR1^f/f^ group. (**Fig. 3A**); however, no changes were observed in females (**Fig. 3B**). Previous reports have demonstrated that extra-synaptic upregulation of NMDAR1/NMDAR2B containing subunits occurs in chronic pain (Hardingham and Bading, 2010; Meng and Shen, 2022; Petralia, 2012). To investigate this further, we isolated synaptic fractions and reexamined NMDAR1 expression in male TNFR1^f/f^ and NexCre^ERT2^::TNFR1^f/f^ mice and determined there was <u>no change</u> in synaptic NMDAR1 receptor subunits (**supplemental Fig. 1**). However, NMDA receptors containing NMDAR2B subunits remained elevated following CCI in both male and female TNFR1^f/f^ and NexCre^ERT2^::TNFR1^f/f^ mice (**Fig. 3C, D)**. To determine if NMDAR1 expression was elevated following CCI in males due to Nex+ neuronal TNFR1 signaling specifically via p38MAPK activation, we quantified expression in NexCre^ERT2^::p38MAPK^f/f^ mice. Following CCI, in males we observed a similar increase in NMDAR1 expression in p38MAPKf/f controls and decreased expression in NexCre^ERT2^::p38MAPK^f/f^ males (**Fig. 3E**). Again, no changes were observed in females (**Fig. 3F**). These data suggest expression of NMDAR1 containing receptors are regulated through a sTNF/TNFR1 signaling nexus in a sex-dependent manner.

**Figure 3.**
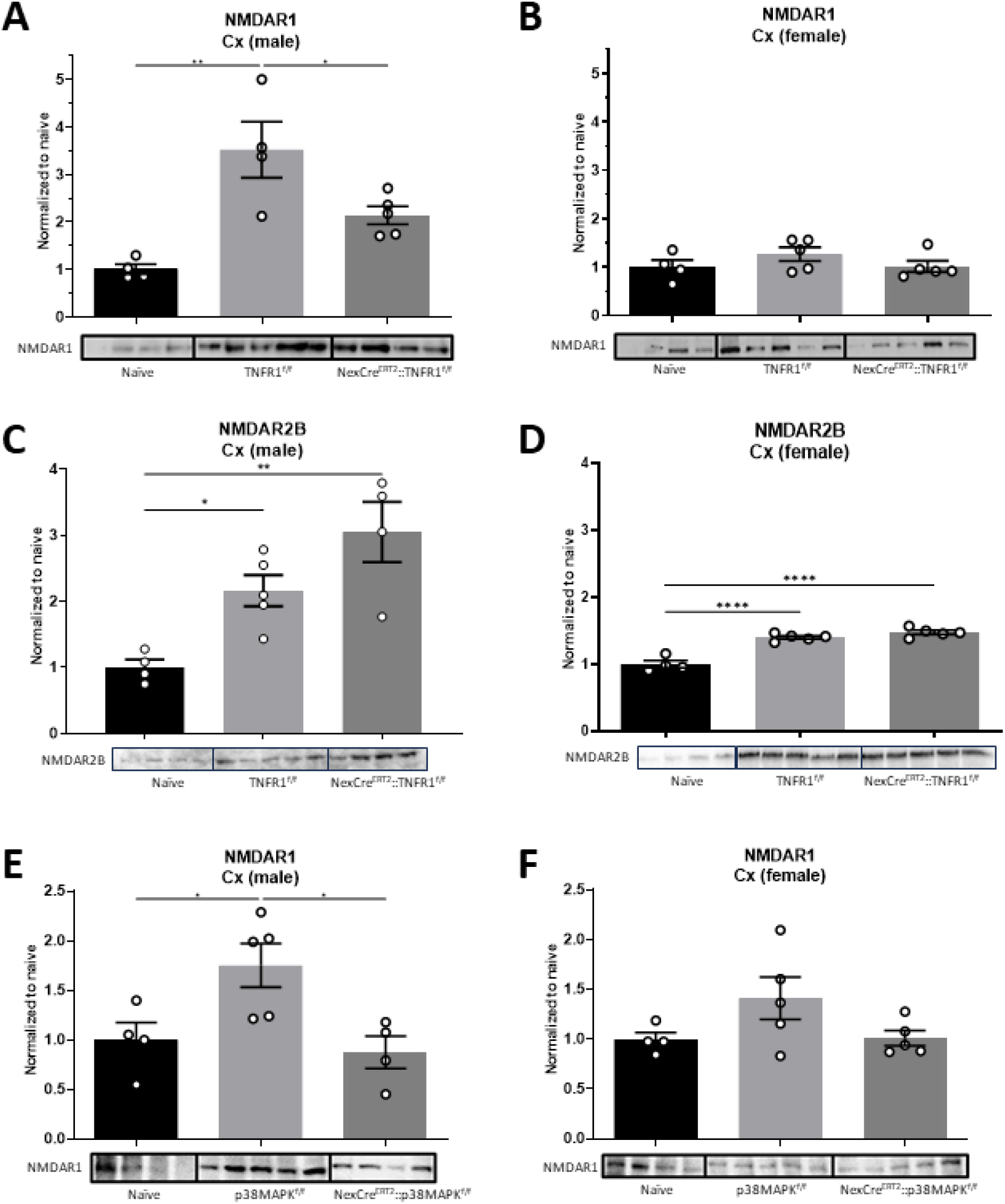
Elevated extrasynaptic NMDAR1 expression in the male cortex is attenuated following selective knockout of TNFR1 or p38αMAPK on excitatory projection neurons. (A) Extrasynaptic NMDAR1 cortical expression following injury is reduced in males following TNFR1 knockout on Nex+ neurons. (B) No differences in NMDAR1 expression compared to controls are observed in females. (C,D) NMDAR2B cortical expression remains elevated in NexCre^ERT2^::p38αMAPK^f/f^ males and females. (E,F) NMDAR1 cortical expression that is normally elevated post-CCI is attenuated in NexCre^ERT2^::p38αMAPK^f/f^ males, but no changes were observed in females (*p<0.05, **p<0.01, *** p<0.001, **** p<0.0001; Data represent mean ± SEM).

### 3.4 The role of ER β in female-typical neuropathic pain

The data presented above (**Figs. 1-3**) and our previously published work demonstrate sex-specific effects of inhibiting sTNF/TNFR1 and p38MAPK on maladaptive plasticity and CNP (Del Rivero et al., 2019). Therefore, we sought to interrogate which estrogen receptors (e.g., ERα and/or ERβ) contribute to the sex-specific effects of sTNF/TNFR1-p38MAPK signaling in CNP. In these studies, we used Xpro1595, a sTNF inhibitor, with either Faslodex (aka Fulvestrant) or PHTPP. Faslodex is an inhibitor of both ERα and ERβ used clinically to treat breast cancer and other disorders (Peekhaus et al., 2004; Rocca et al., 2018). PHTPP, on the other hand, is a highly selective ERβ antagonist (e.g., silent antagonist), with ∼36-fold selectivity for ERβ over ERα (Compton et al., 2004). With these inhibitors, we sought to determine the relationship between estrogen and therapeutic efficacy of sTNF inhibition. Following injury, female mice were treated with vehicle, Faslodex, Xpro1595, or combinations of Faslodex and Xpro1595 as previously described (Del Rivero et al., 2019). As shown in **Fig 4A**, combination therapy of Faslodex at the time of CCI and Xpro1595 one-week post-CCI significantly reduced CNP in females whereas either drug alone did not. Next, we investigated the therapeutic potential of combining PHTPP with Xpro1595 or SB203580, a potent p38MAPK inhibitor. Combination therapies of PHTPP with either Xpro1595 (**Fig. 4B**) or SB203580 (**Fig. 5A**) were highly therapeutic for CNP in females, whereas individual drugs were not (**Figs. 4B,5A**). Interestingly, combination therapies of SB203580 and PHTPP in females were equally as effective as SB203580 alone in males (**Fig. 5A**). These, and other data, suggest that inhibition of ERβ signaling renders females “male-like” with respect to CNP. Finally, we demonstrated that female cortical ERβ expression after CCI was similar to that of naïve levels when treated with PHTPP (**Fig. 4C**). However, spinal ERβ expression remained unchanged in all groups (**Fig. 4D**). Following PHTPP administration, cortical p38αMAPK activation appears to be reduced compared to vehicle-treated controls; although, there was no statistical significance (**Fig. 5B**). No changes were observed in the lumbar spinal cord (**Fig. 5C**). Mechanisms through which PHTPP may modulate ERβ expression have yet to be determined.

**Figure 4.**
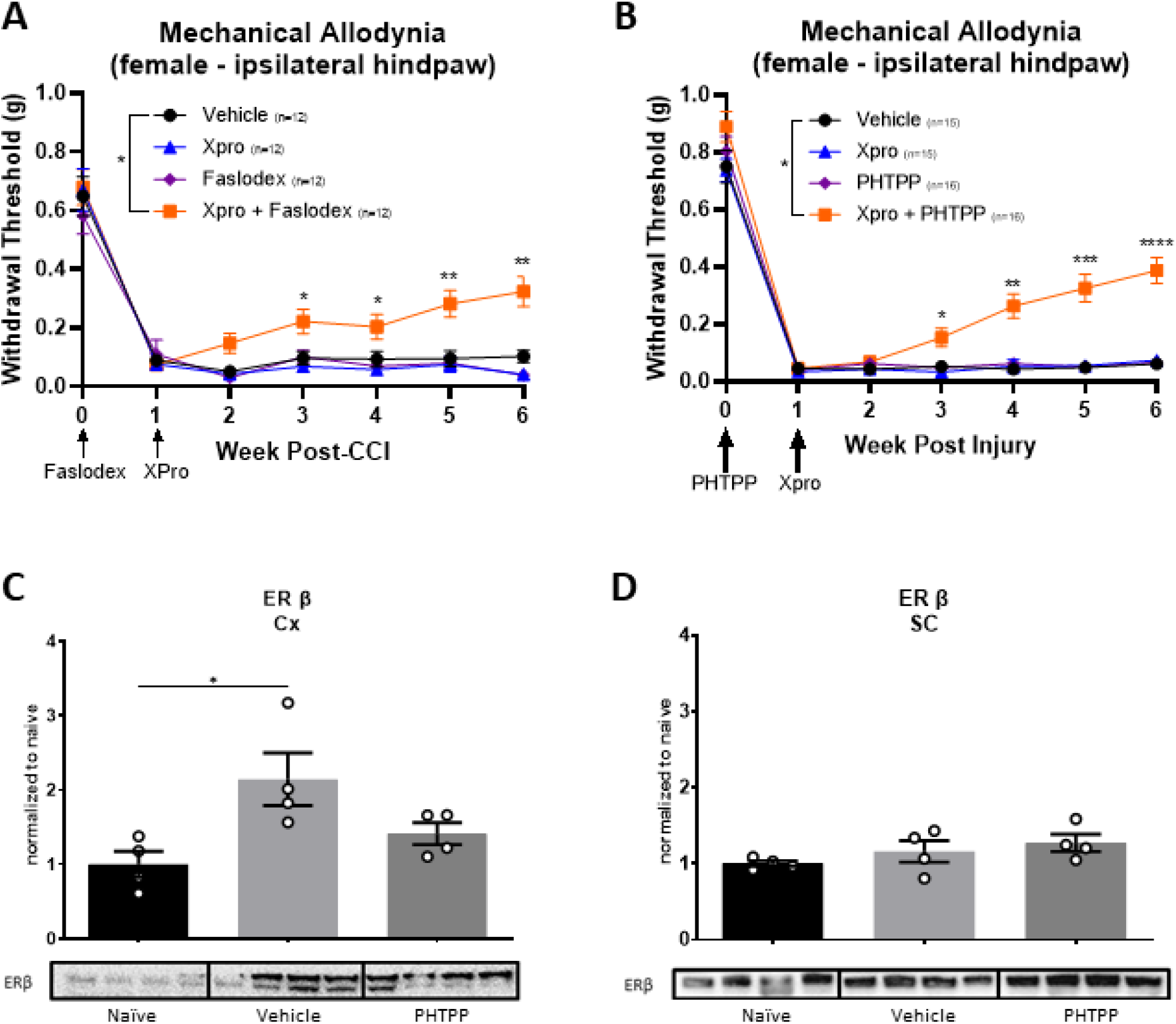
Inhibition of soluble TNF and estrogen receptors, particularly ER β, reduces mechanical allodynia in females. (A,B) Xpro paired with Faslodex, a general ER inhibitor, or PHTPP, an ER β inhibitor attenuates withdrawal threshold. (C) Cortical ER β is elevated following CCI; however, following treatment with PHTPP there is no significant difference compared to naïve mice. (D) ER β expression in the lumbar cord remains unchanged. (E) Representative images taken from anterior cingulate cortex demonstrates reduced ER β expression following treatment with PHTPP [ERβ expression = 1^st^ column; NeuN = 2^nd^ column; Merged = 3^rd^ column] (*p<0.05, **p<0.01, *** p<0.001, **** p<0.0001; Data represent mean ± SEM).

**Figure 5.**
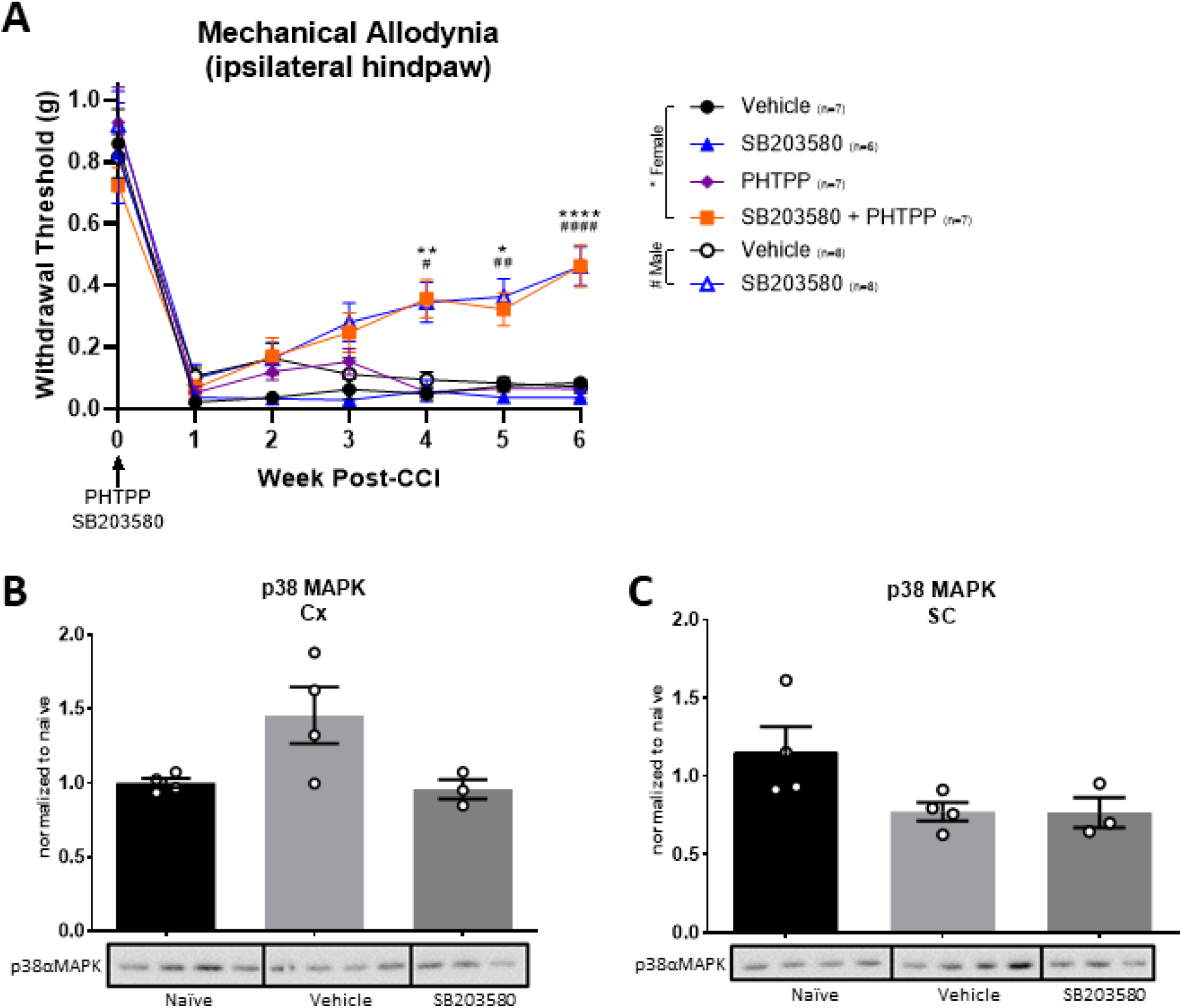
In females, combination therapies of PHTTP and SB2030580 is therapeutic for CNP but individual therapies are not. (A) SB203580 alone increases withdrawal threshold in males post-CCI, but females require SB203580 and PHTPP to produce similar alleviation. (B) Although not significant, it appears that cortical p38αMAPK is elevated in vehicle treated groups and is reduced to naïve-like activation levels following SB203580 treatment in females. (C) However, vehicle- and SB203580-treated groups express comparable p38αMAPK levels in the lumbar cord (*p<0.05, **p<0.01, *** p<0.001, **** p<0.0001; Data represent mean ± SEM).

## 4. DISCUSSION

The contribution of biochemical sex differences to the increased prevalence of CNP amongst females and males and the role of sex hormones in relation to pain tolerance has been well established (Claiborne et al., 2006; Li et al., 2009). Crucial mediators of CNP include neuroinflammation and maladaptive plasticity; however, we have yet to elucidate the intersect between these critical mediators and sex differences, such as estrogen. We demonstrate that CNP in males, but not females, is dependent on TNFR1 signaling and p38αMAPK activation in Nex+ supraspinal excitatory neurons. Further, inhibition of ER β renders females “male-like” with respect to inhibiting sTNF and p38MAPK as viable therapeutic targets. Altogether, these data indicate the importance of supraspinal circuitry in chronification of pain and elucidate potential mechanisms through which male and female pain may be treated.

### 4.1 TNFR1 and p38**α**MAPK signaling in supraspinal excitatory projection neurons mediate CNP in a sex-specific manner

The link between neuroinflammation and persistent neuronal hyperexcitation and their contribution to both the induction and maintenance of CNP is well established. Previously, we have demonstrated that neuroinflammatory mechanisms of male CNP are heavily mediated by TNFR1 signaling (Del Rivero et al., 2019). TNFR1 signaling is known to activate apoptotic and pro-inflammatory pathways that include NF-κB and p38MAPK (Jaco et al., 2017; Wajant and Siegmund, 2019). Both pathways exacerbate inflammation and chronic glial activation that then furthers the production of cytokines and promotes increased neuronal activity (Ji et al., 2018).

Previous research has demonstrated that supraspinal signaling in cortical and hippocampal excitatory projection neurons is critical for the chronification of pain in males (Kelly et al., 2016; Shiers and Price, 2019). In the brain, pyramidal neurons comprise most excitatory neurons in the cortex, hippocampus and amygdala and heavily employ NMDAR pathways (Spruston, 2008; Wang et al., 2018). Nex+ neurons are a subset of these pyramidal neurons located supraspinally. Excitatory pathways associated with male CNP may, in part, be mediated via the TNFR1-p38αMAPK pathway where sTNF leads to increased presence of extrasynaptic NMDARs as a consequence Woods et al., 2021). In several disease and injury paradigms, extrasynaptic NMDARs are linked to cell death through pro-apoptotic pathways and are activated by glutamate via synaptic spillover or ectopic release (Hardingham and Bading, 2010; Petralia, 2012). When TNFR1 and p38αMAPK are knocked out on Nex+ neurons, cortical NMDAR1 extrasynaptic expression is reduced solely in males. However, extrasynaptic NMDAR1 expression is found in both neuronal and glial populations aiding in neuron-to-glia communication (Dzamba et al., 2013; Hardingham and Bading, 2010; Lalo et al., 2006). Future studies are needed to determine whether the aforementioned changes in NMDAR1 are glial or neuronal in their extrasynaptic presence.

Microglial activity via p38 signaling was previously demonstrated by others to preferentially contribute to neuropathic pain in male mice (Paige et al., 2018). However, p38MAPK signaling is not restricted to spinal microglia and occurs in neuronal/non-neuronal populations located supraspinally and within dorsal root ganglion (Crown et al., 2008; Jin et al., 2003). These studies employed the use of intrathecal skepinone, a p38MAPK selective inhibitor, which may also inhibit supraspinal, spinal, and DRG neuronal/non-neuronal p38MAPK signaling (Paige et al., 2018)(56)(Paige et al., 2018). Our studies used a powerful genetic approach to determine that neuronal p38αMAPK signaling in neuropathic pain plays a critical role in pain chronification. While previous studies demonstrate phosphorylation of p38MAPK is more frequently associated with sTNF-associated microglial inflammation, our data support that supraspinal neuronal p38αMAPK is also a critical regulator of CNP following inflammation via peripheral injury.

### 4.2 ERβ regulates the behavioral phenotype of female CNP

Chronic pain is more prevalent among females, and this is observed across a spectrum of conditions (Bartley and Fillingim, 2013; Casale et al., 2021; Unruh, 1996). Mounting evidence exists that shows differential immune system response between females and males during pain, which may account for differences in immune targeted therapeutic efficacy between sexes (Gregus et al., 2021; Rosen et al., 2017). Estrogen receptors (ERs) are widely distributed among nociceptive regions of the CNS, including the thalamus and anterior cingulate cortex, and are known to modulate sensory processing and transduction (Chen et al., 2012, 2021; Li et al., 2009; Shughrue et al., 1997; Zhang et al., 2020). Specifically, estrogen and ERs are known to contribute to supraspinal neuronal maladaptive plasticity associated with chronic pain and modulate immune cell response (Chen et al., 2021; Khan and Ansar Ahmed, 2016; Sun et al., 2019). We have previously demonstrated that following CCI, pharmacological and genetic inhibition of TNFR1 signaling is selectively therapeutic in males (Del Rivero et al., 2019). However, in ovariectomized females, inhibition of TNFR1 signaling has “male-like” therapeutic efficacy. Our findings suggest that ER β interferes with therapeutic efficacy via TNFR1 signaling inhibition. More specifically, combination therapy in females with ER β, sTNF, and p38MAPK inhibitors rendered them more “male-like” with regards to recovery from allodynia. Previous work has shown that ER β activation in cortical neurons increases the density of dendritic spines as well as clustering of postsynaptic density-95 at the membrane (Srivastava et al., 2010). Together, these data suggest that cortical ER β may modulate post-synaptic excitatory signaling and, in turn, enhance glutamate-mediated excitotoxicity. Inhibition of sTNF/TNFR1 signaling pathways reduces excitatory signaling; however, ER β may enhance cortical excitotoxicity. By blocking ER β in females, we reduce this enhanced excitotoxicity and allow females to be more amenable to anti-TNF therapies. Future studies are needed to determine efficacy for other pain categories such as thermal and spontaneous pain.

## 5. CONCLUSIONS

In summary, our data demonstrate that supraspinal neuronal TNFR1/p38αMAPK signaling is critical in chronification of pain with male specificity. When TNFR1/p38αMAPK signaling is selectively knocked out on supraspinal excitatory neurons, the transition from acute to chronic pain is avoided in males. Furthermore, when ER β is inhibited in females and sTNF and p38MAPK inhibitors are administered, pain does not transition from an acute to chronic state. With these data we propose that supraspinal neuronal TNFR1/p38αMAPK signaling is critical to pain chronification in a sex-specific manner but poses a novel therapeutic route for treatment of CNP.

## Key Words/Abbreviations

(sTNF): Soluble tumor necrosis factor
(TNFR1): TNF receptor 1
supraspinal excitatory projection neurons
(p38MAPK): p38 mitogen-activated protein kinase
(ER β): estrogen receptor beta
(CNP): chronic neuropathic pain

---

**Supplemental Figure 1.**
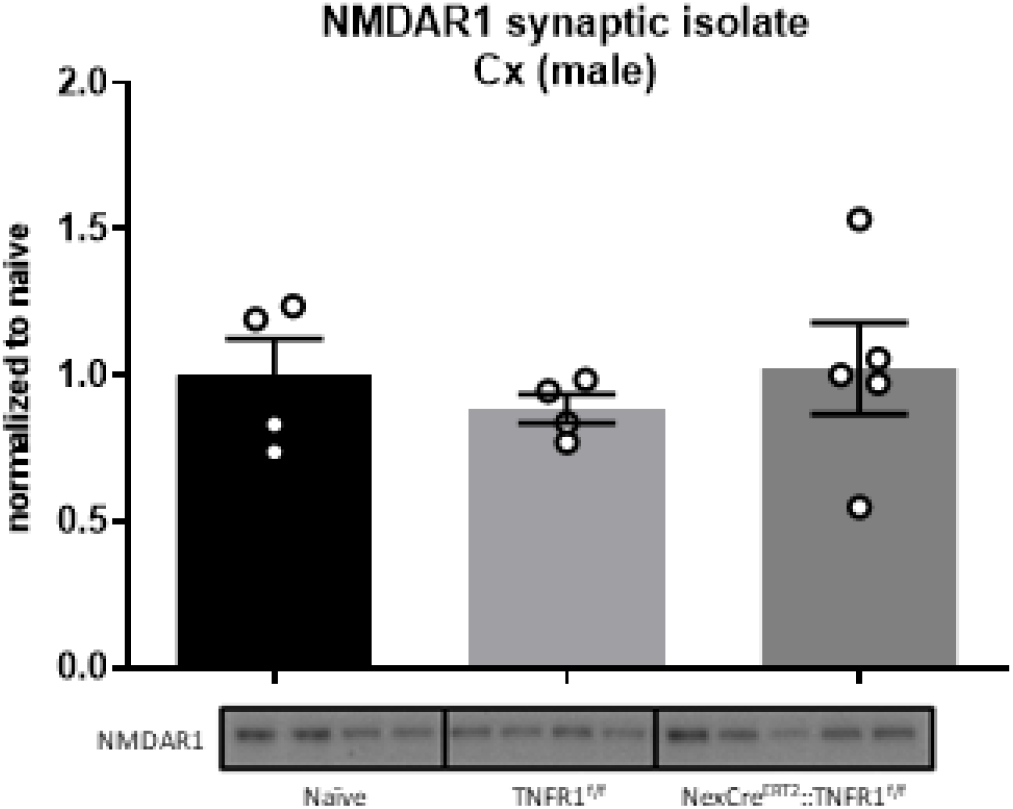
There are no changes in male cortical expression of synaptic NMDAR1 (Data represent mean ± SEM)

**Supplemental Figure 2.**
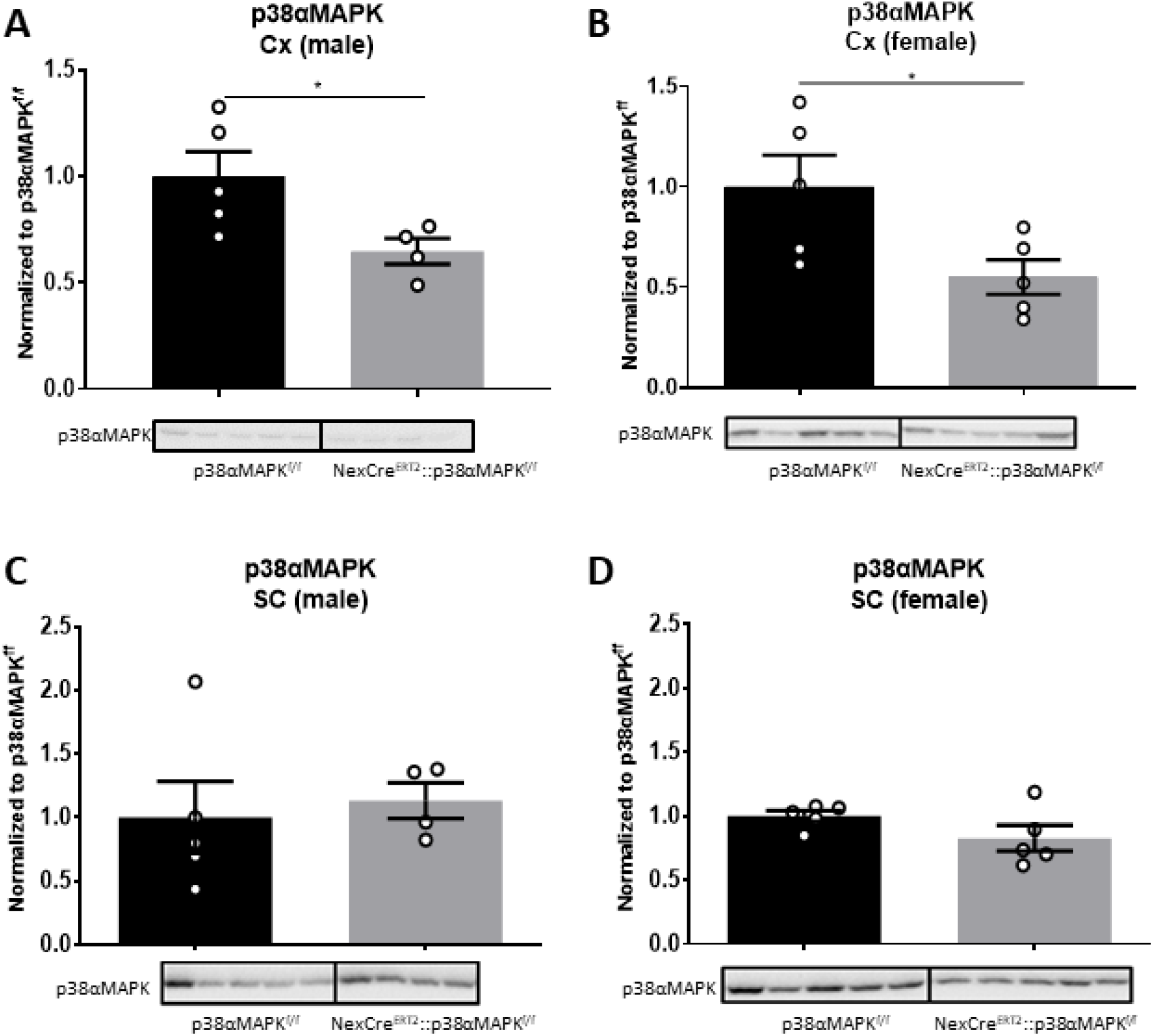
Cortical p38αMAPK is reduced in both female and male NexCreERT2::p38αMAPK mice. (A,B) p38αMAPK expression in the cortex is reduced in both female and male NexCreERT2::p38αMAPK mice. (C,D) However, there are no changes in spinal p38αMAPK for both sexes (*p<0.05; Data represent mean ± SEM).

